# Advancing Biogeographical Ancestry Predictions Through Machine Learning

**DOI:** 10.1101/2025.01.31.635898

**Authors:** Carola Sophia Heinzel, Lennart Purucker, Frank Hutter, Peter Pfaffelhuber

## Abstract

Tools like Snipper or the Admixture Model count as state-of-the-art methods in forensic science for biogeographical ancestry. However, they have not been systematically compared to classifiers widely used in other disciplines. Noting that genetic data have a tabular form, this study addresses this gap by benchmarking forensic classifiers against TabPFN, a cutting-edge, general-purpose machine learning classifier for tabular data. The comparison evaluates performance using metrics such as accuracy—the proportion of correct classifications—and ROC AUC. We examine classification tasks for individuals at both the intracontinental and continental levels, based on a published dataset for training and testing. Our results reveal significant performance differences between methods, with TabPFN consistently achieving the best results for accuracy, ROC AUC and log loss. E.g., for accuracy, TabPFN improves SNIPPER from 84% to 93% on a continental scale using eight populations, and from 43% to 48% for inter-European classification with ten populations.

## 1 Introduction

Classification of individuals into ancestral populations, i.e., inferring biogeographical ancestry (BGA) from DNA traces, is a fundamental task in forensic science (Wen et al., 2023). This process is particularly useful for identifying disaster victims (Phillips et al., 2009) and in broader forensic investigations (Phillips et al., 2007b). Additional applications include detecting population stratification (Dang et al., 2005; Min et al., 2000; Gan et al., 2003). Researchers commonly classify individuals into several continental populations such as Africa, Europe, East Asia, South Asia, Admixed Americans, and Oceania (Cheung et al., 2017; Wang et al., 2007), or on an intracontinental level.

The statistical task of classification can be broken down into two steps. First, genetic markers have to be chosen which are able to distinguish between populations, i.e. with frequency differentials between populations. Second, these Ancestry Informative Markers (or AIMs) are used as features in a classification task. Here, various algorithms are used, leveraging genetic distance, regression, or Bayesian inference (Cheung et al., 2017). Tools like the naive Bayes classifier Snipper (Phillips et al., 2014) have been used alongside multinomial logistic regression (McNevin et al., 2013) and the Genetic Distance Algorithm (Frudakis et al., 2003). Furthermore, the Admixture Model, as implemented in tools like Structure (Pritchard et al., 2000) or Admixture (Alexander et al., 2009), is widely used for classification (Cheung et al., 2017). More recently, other machine learning methods such as XGBoost (Alladio et al., 2022; Šorgić et al., 2025; Kloska et al., 2023), Partial Least Squares-Discriminant Analysis (PLS-DA) (Alladio et al., 2022), and random forests (Kloska et al., 2023; Zaorska et al., 2019; Song et al., 2019) have been applied in forensic genetics to ancestry prediction.

Several challenges must be taken into account for BGA. First, classification becomes increasingly challenging for more genetically similar populations, which are frequently found within continents. Second, admixed individuals, such as Americans, present difficulties in classification (Lao et al., 2010; Llamas et al., 2016). Third, marker selection significantly affects classification quality (Cheung et al., 2017), with ideal markers showing large differences in allele frequencies between populations (Rosenberg et al., 2003). Fourth, selecting populations as classes must lead to results which are both, reliable and helpful in further investigations.

The choice of Ancestry Informative Markers (AIMs) is frequently performed by finding alleles with large frequency differentials between populations using expert knowledge (Kidd et al., 2014; Phillips et al., 2014; Ruiz-Ramírez et al., 2023). However, feature selection for classification is also a well-known task in machine learning, which has been applied to the choice of AIMs (Wathen et al., 2019; Pfaffelhuber et al., 2020b). The goal is to find a set which leads to a reduced sequencing effort, focusing on 50–200 informative markers (relative to the classes under study) instead of thousands (Ko et al., 2023). However, Heinzel et al. (2024) showed that marker quality criteria extend beyond allele frequency differences. Additionally, forensic contexts often face challenges with missing data (Cheung et al., 2017).

Cheung et al. (Cheung et al., 2017) compared several classification approaches, including Snipper (Phillips et al., 2014), multinomial logistic regression (McNevin et al., 2013), and the Genetic Distance Algorithm (Frudakis et al., 2003). They also analyzed the Admixture Model, concluding that “*STRUCTURE was the most accurate classifier for both complete and partial genotypes in non-admixed individuals across four reference populations, including populations with suspected admixture*.*”* This study is complemented by a comparison to XGboost and PLS-DA in (Alladio et al., 2022), who conclude that such machine learning tools have a higher accuracy when compared to the above methods. We complement this study by using TabPFN, a new foundational model which was shown to outperform existing models for tabular data (Hollmann et al., 2025).

Ideally, a classifier predicts the population of every individual with 100% ancestry of one population correctly with prediction probability 1. Additionally, according to Cheung et al. (2017), for admixed individuals the prediction probability should be equal to the ancestry proportions of the genome. Several measures of non-optimal classifiers, such as the misclassification rate, the logloss (sum of logs of misclassification probabilities), accuracy (fraction of correctly classified samples), the ROC AUC (the area under a curve of false positives against true positives) are among the most common statistics.

Machine learning has already seen applications in forensic science, including haplogroup prediction from Y-STR data (Swaminathan et al., 2015), estimation of the number of contributors to a trace (Bouakaze et al., 2020), and prediction of the age or post-mortem interval (Barash et al., 2024). While machine learning methods have been employed to predict physical appearance from DNA (Katsara et al., 2021) as well as BGA (Wathen et al., 2019), the application of such methods have been criticized for their lack of fit to the specific needs in the forensic field (Barash et al., 2024). Our work aims to address this gap by applying state-of-the-art machine learning techniques to the BGA classification problem.

## 2 Material and Methods

### 2.1 Choice of the Data and the Marker Set

There are different widely used marker sets in forensic genetics, (e.g. Phillips et al., 2014; Kidd et al., 2014; Xavier et al., 2020; Ruiz-Ramírez et al., 2023). Throughout our study, we adopt the dataset from Ruiz-Ramírez et al. (2023) (their Supplementary Table S1A), which uses the markers from the VISAGE Enhanced tool (Xavier et al., 2020). On the one hand, the choice of the markers influences the output of the classifiers. On the other hand, “most of the BGA SNPs [in the VISAGE Enhanced Tool] are already well established for forensic use” (Ruiz-Ramírez et al., 2023). So, we choose the VISAGE Enhanced tool, since this is the latest widely used marker set for ancestry prediction. This set of markers has 104 autosomal markers, 29 of which are multiallelic. This data set consists of 2504 individuals from the 1000 genomes project (Consortium et al., 2015), 929 HGDP-CEPH samples (Phillips et al., 2013; Santos et al., 2016), 137 samples from the Middle East (Almarri et al., 2021), 130 samples from the SGDP (Mallick et al., 2016), and 402 samples from the Estonian Biocentre human genome diversity panel (Pagani et al., 2016).

We consider two cases:

1. Continental level combining *‘AFRICAN’,’EAST AFRICAN’* and *‘ADMIXED AFRICAN’* to Africa (AFR), combining *‘MIDDLE EAST’* and *‘(SANGER) MIDDLE EAST’* to Middle East (ME), combining *‘EUROPEAN’* and *‘ROMA’* to Europe (EUR), East Asia (EAS), South-East Asia (SAS), Oceania (OCE), *‘CENTRAL ASIAN’* (CAS), *‘NORTH AFRICAN’* (NAF) and combining *‘ADMIXED AMERICAN’* and *‘AMERICAN’* to Admixed America (AMR). This makes a total of nine classes.
2. Intra-European level, only considering the EUR classes from above with more than 20 individuals, using *(CEPH) with N & W European ancestry’* (CEU), *‘Finnish in Finland’* (FIN),*’Toscani in Italia’* (TSI), *‘Italy - Sardinian’* (SAR),*’British in England and Scotland’* (GBR), *‘Iberian population in Spain’* (IBS), *‘Toscani in Italia’, ‘Russia - Russian’* (RUS), *‘France - French Basque’* (BAS), *‘France - French’* (FRA), and *‘Turkey’* (TUR) as classes. These are a total of ten classes. We did not combine BAS and FRA as Flores-Bello et al. (2021) mentioned that there are big differences between basques and people from France. Similarly, we considered TSI and SAR as different populations as there genetic distance is significant (Chiang et al., 2018).

The number of individuals is shown for both cases in Figure 1, where we see that some classes are under-represented in the dataset.

**Figure 1.**
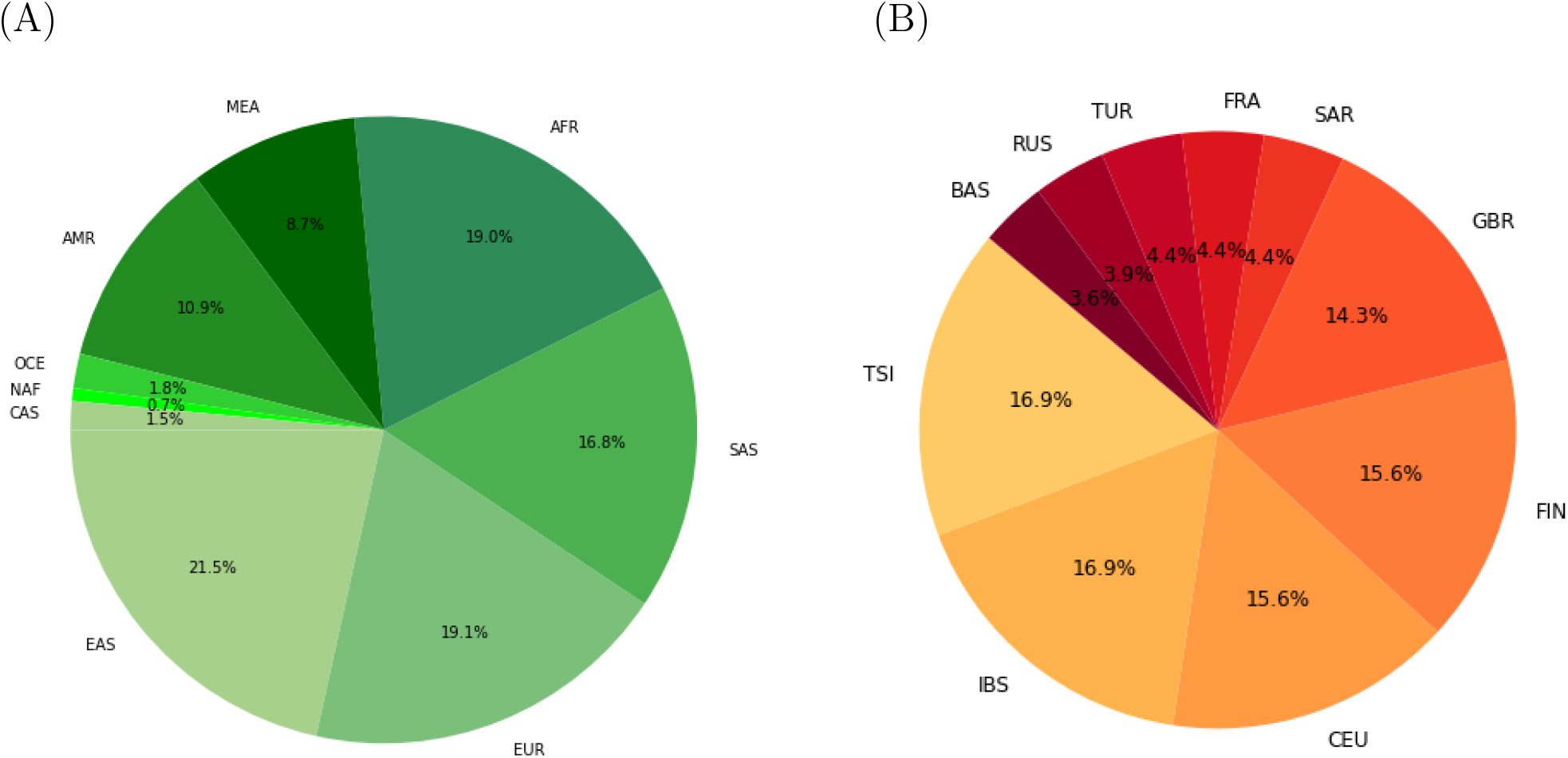
(A) Overview of the number of individuals per population in case (1), i.e. for continental classification. In this case, there are a total of 4342 individuals. (B) Overview of the number of individuals per population for case (2), i.e. intra-European classification. In this case, there are a total of 635 individuals. Total sample sizes for all populations can be found in Tables S1 and S2 in the Supplement.

### 2.2 Classification

For classification, we compare the following methods: (i) Snipper, which is a version of a naive Bayes classifier (Phillips et al., 2007a); (ii) the Admixture Model (AM) (in its supervised setting), which assumes that every allele in individual *i* has a chance of *q*_*ik*_ to come from population *k* (Pritchard et al., 2000; *Alexander et al., 2009)*, and individual *i* is classified into population *k* if *q*_*ik*_ is maximal; (iii) PLS-DA which has already been used by (Alladio et al., 2022) in the forensic context; and (iv) TabPFN, a novel foundational model for tabular data (such as genetic data) based on a neural network (Hollmann et al., 2025), (v) XGBoost (Chen and Guestrin, 2016) and (vi) Random Forest (Breiman, 2001). Note that the model-based methods (i) and (ii) treat markers as independent, while the machine-learning (black-box) methods (iii), (iv), (v) and (vi) incorporate joint segregation of markers.

Other classifiers, such as gradient-boosted decision trees (Chen and Guestrin, 2016; Ke et al., 2017; Prokhorenkova et al., 2018), or logistic regression (Cheung et al., 2019), would as well be applicable to genetic (i.e., tabular) data. However, Hollmann et al. (2025) have shown that TabPFN dominantly outperforms other classifiers, including those mentioned, for small tabular datasets. We evaluate if these finding also holds for ancestry prediction.

Neural networks play a central role in machine learning today. While convolutional neural networks have become the foundation for models for image (and video) data, the transformer neural network architecture is the foundation for large language models (such as ChatGPT). With TabPFN, a foundation model for tabular data was introduced, making it possible to apply transformer neural networks to any classification problem on tables instead of images or text. In detail, TabPFN is a transformer (Vaswani et al., 2017) neural network that was pretrained on millions of synthetic datasets to learn how to solve classification tasks (Hollmann et al., 2022, 2025). This pre-trained neural network can then solve new classifications for real-world datasets in one forward pass through the network. We stress that TabPFN is a pre-trained neural network, and the weights (which are independent of the application context) are open and free to use^1^.TabPFN excels in handling small to medium-sized datasets with up to 10,000 samples and 500 features.

TabPFN has a higher usage cost than traditional classifiers because it needs a graphics processing unit (GPU) to predict efficiently. However, TabPFN is a relatively small model (as compared to large language models), since it contains millions of parameters rather than billions (or even trillions). As a result, it can run efficiently on consumer-grade hardware, such as desktop GPUs or integrated GPUs (e.g., Apple’s M-series chips), with about 8GB of VRAM. Despite the usage cost, we select TabPFN for our comparison because it requires substantially less training time if a GPU is available (Hollmann et al., 2025).

We evaluate each classifier by training the model on a subset of the data and evaluating its prediction on a holdout set. This experiment is repeated 50 times by using 10-repeated 5-fold cross-validation with stratification from scikit-learn (Pedregosa et al., 2011). We score predictions using accuracy, log loss and ROC AUC^2^, following best practices from machine learning (Provost et al., 1998; McElfresh et al., 2023; Gijsbers et al., 2024; Vovk, 2015). ROC AUC measures the ability to distinguish between classes and 1, indicating perfect discrimination, and 0.5, representing random guessing. Analogously to previous evaluation metrics in forensics (Pfaffelhuber et al., 2020a; Resutik et al., 2023), we also count the number of true positive and false negative samples to compute a confusion matrix. More precisely, for every of our 50 repetitions, we sum up the number of individuals from class *i*, e.g., Europe, that are classified into population *j*, e.g., Africa. The resulting percentiles are depicted as a confusion matrix.

### 2.3 Code availability

All code for our analysis can be downloaded from GitHub. On this website you can also find an overview of the hyperparameter that we used and the versions of the Python packages. We include mostly Python scripts for selecting individuals from the dataset, and a locally running web-interface for easier use of classification. We also set up this interface online.

## 3 Results

First, we compare the evaluation metrics ROC AUC, Accuracy and Log Loss between the different classifiers and between the continental and intra-European classification. Second, we compare the corresponding confusion matrices of the classification methods to each other. We provide results for all combinations of classifier and continental and inter-European classification.

### 3.1 Evaluation Metrics

The ROC AUC (between 0.5 and 1, higher is better), Accuracy (between 0 and 1, higher is better) and the Log loss (greater than 0, lower is better) of the classification in case (1) and (2) can be found in Figure 2 (numerical values in Tables S3, S4, S5 in the Supplement).

**Figure 2.**
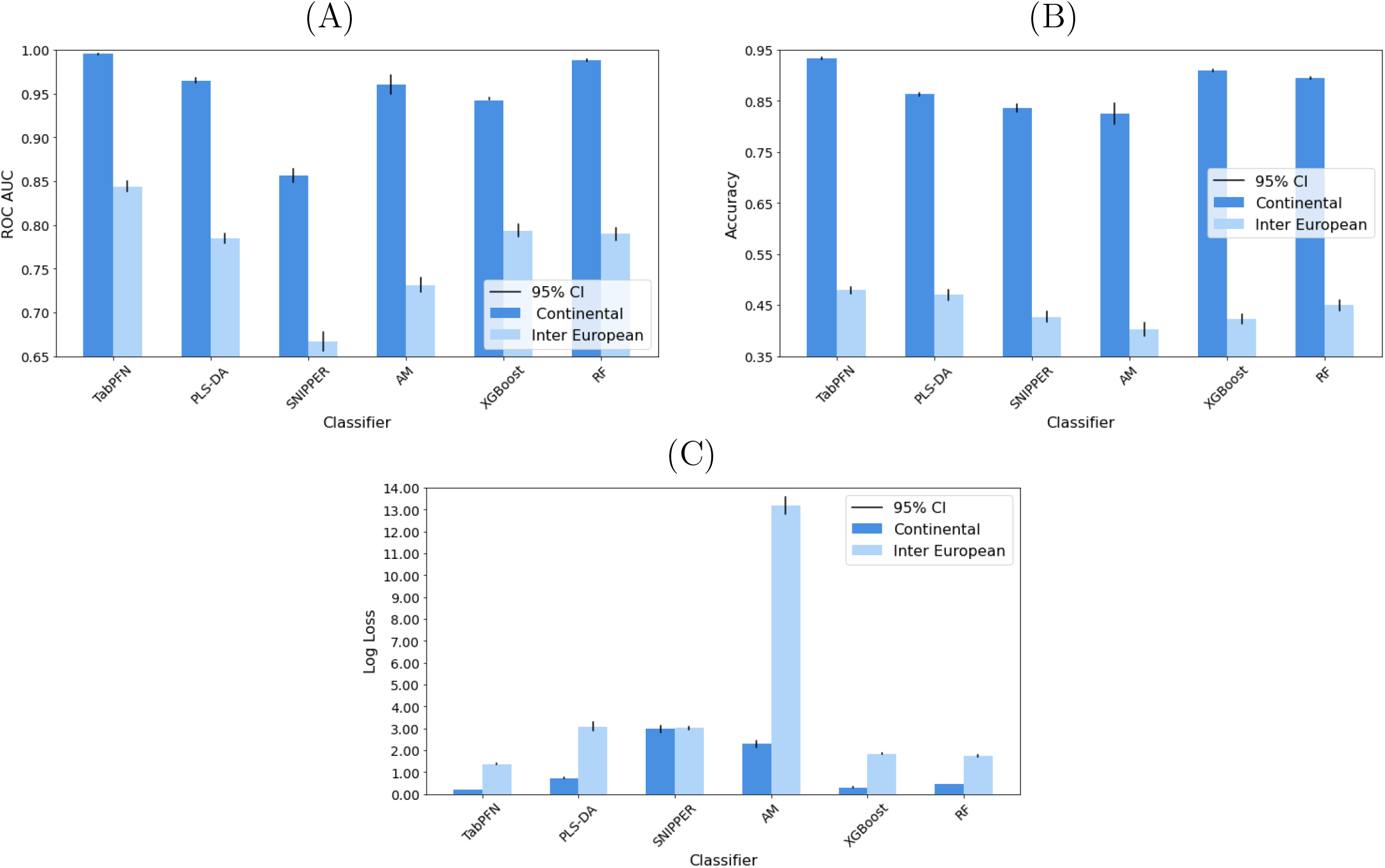
Evaluation metrics for continental (1) and inter-European (2) classification. The black line represents the 95% confidence interval. (A) ROC AUC; (B) Accuracy; (C) Log Loss. AM stands for Admixture Model and RF stands for Random Forest. For exact numbers, see Tables S3, S4 and S5.

For all three evaluation metrics TabPFN is the best classification method in both, continental and inter-European classification. While performing worse than TabPFN, PLS-DA is better than Snipper and the Admixture Model in most cases. ROC AUC for Snipper is worse than for Admixture Model while the accuracy and the log loss of Snipper are better than for the Admixture Model. Altogether, all four considered classification methods work well for continental classification. Unsurprisingly, performance drops significantly for inter-European classification. Even TabPFN only classifies 48% of samples correctly, which can be explained by the relatively large number of classes (10) and the high similiarity between the classes.

### 3.2 Confusion Matrices

For more precise results which distinctions work well and which don’t, we display confusion matrices for all eight cases. (Cases (1) and (2), six classifiers.) The results are shown in Figures 3 and 4 for TabPFN, Snipper, DLS-PA and Admixture, and in Figure S1 in the supplement for XGBoost and Random Forest.

**Figure 3.**
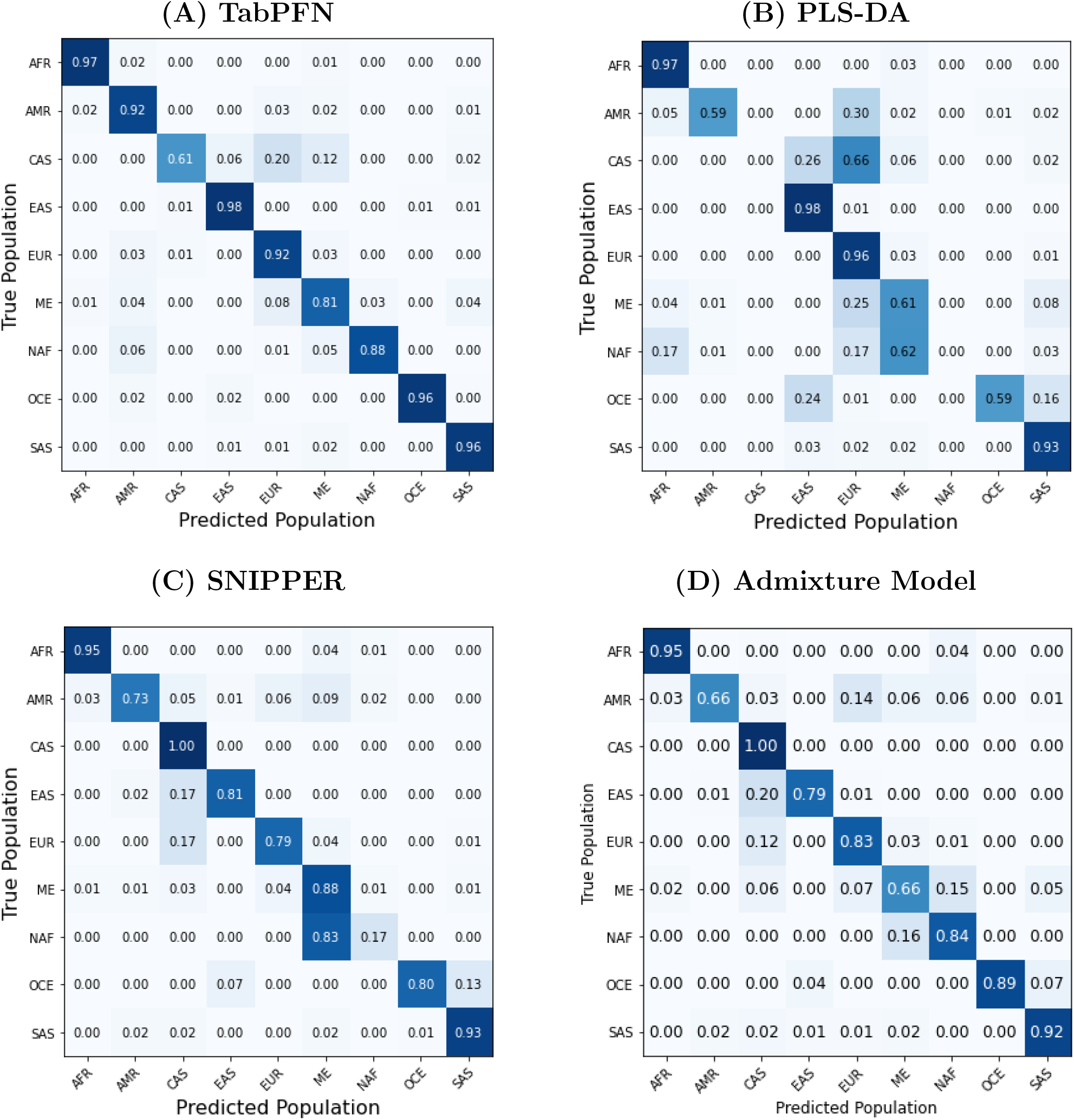
Confusion matrices for continental classification (case 1). All rows sum to 1 up to a rounding error. **(A)** TabPFN; **(B)** PLS-DA; **(C)** SNIPPER; **(D)** Admixture Model.

**Figure 4.**
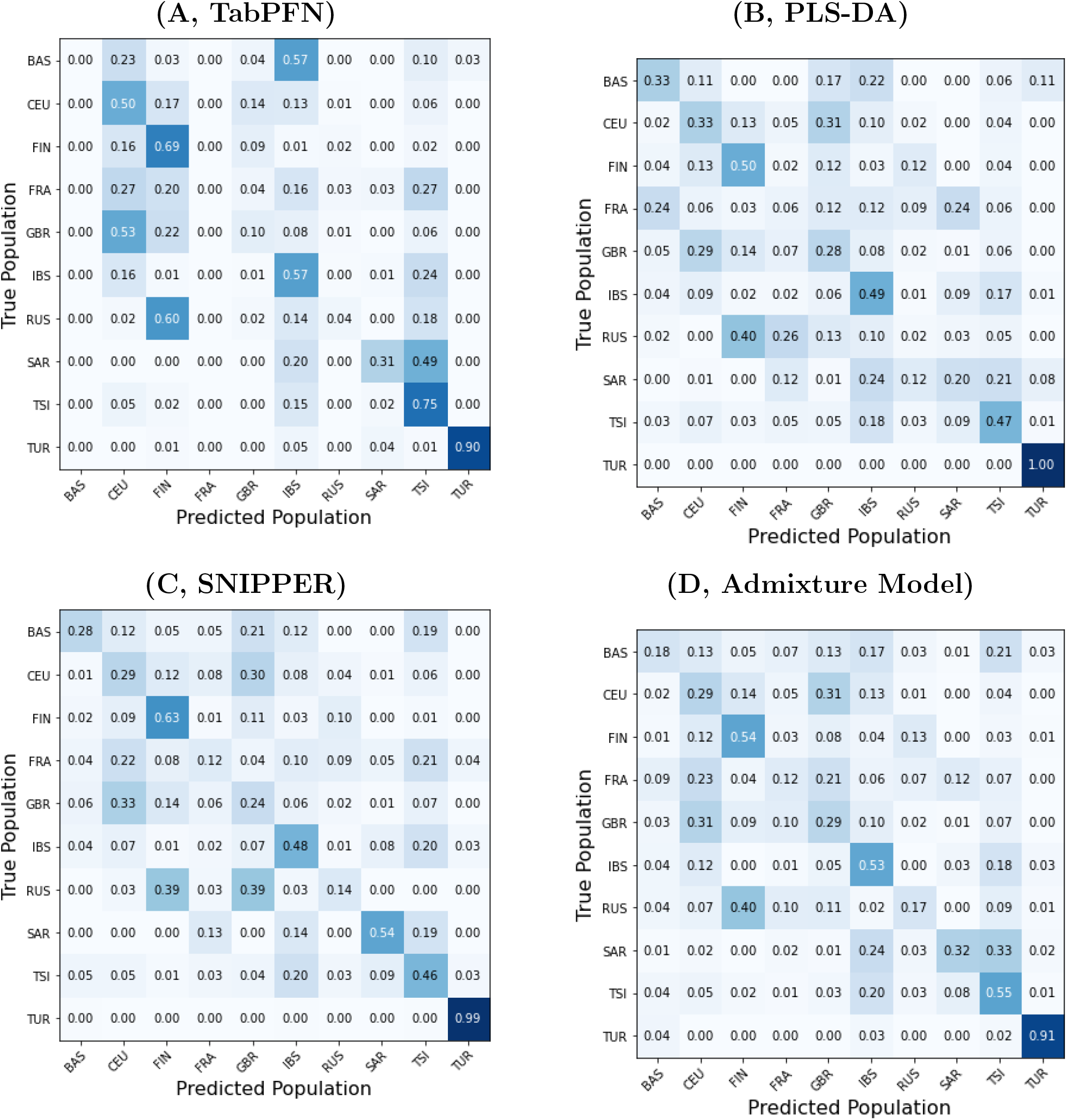
Confusion matrices for intra-European classification (case 2). All rows sum to 1 up to a rounding error. (A) TabPFN; (B) PLS-DA; (C) Snipper; (D) Admixture Model

For continental classification, we see that the classification works very well for all populations except for ME and NAF in all four methods. In 81% of the considered cases, TabPFN classifies ME individuals correctly, which is only outperformed by Snipper (88%). However, Snipper misclassifies the majority of NAF into ME (only 17% being correct), whereas TabPFN achieves 88% also in NAF. PLS-DA and the Admxiture Model are similar to Snipper in the confusion between ME and NAF. Another interesting population is Central Asian. (Note that this data is from EGDP (Pagani et al., 2016) and only contains 64 samples.) Here, TabPFN only achieves 61%, whereas methods which treat markers independently (Snipper, Admixture Model) achieve 100%.

In general, intra-European classification (case (2)) has a higher error rate than continental classification, as can be seen from the confusion matrices in Figure 4 (and Appendix C). With TabPFN, we see that people from GBR are often (in 53% of the cases considered) classified into CEU. Furthermore, people from CEU are often classified as GBR (in 14% of the cases considered). The number of mis-classifications for people from SAR is in general very high and the number of miss-classifications of RUS into FIN is also large.

In comparison to TabPFN, the percentage of correct classifications of people from some populations is even higher for Snipper or for the Admixture Model. An example for this are the individuals from TUR with Snipper and the ones from SAR with the Admixture Model.

## 4 Discussion

In our study, we test six classifiers on well-established (training and test) data, which was evaluated in the VISAGE project (Xavier et al., 2022). We only use classifiers which determine their own uncertainty by reporting probabilities for class membership. Our results show that TabPFN produces the highest average ROC AUC, the highest average accuracy, and the best average log loss for our data. Especially when the populations are exclusively from Europe, TabPFN strongly outperforms all other methods, although the mis-classification rate is still high (52%).

Like any black-box classifiers that is not informed about which variables are sensitive, TabPFN purely learns the statistical patterns in the training data, even if these are based on biases of human curators. For an extension of TabPFN that alleviates such biases with counterfactual fairness, please refer to Robertson et al. (2024). Moreover, the number of individuals per population should be chosen carefully because a small number does not provide enough information for learning, reducing the accuracy of the classifier. As for other black-box classifiers, we generally do not have a guarantee that TabPFN performs well on any real-world data, and TabPFN’s performance should always be validated. However, TabPFN already excels for a broad range of tabular data (Hollmann et al., 2025), and the data from forensic genetics we investigated in this study. Unlike other classifiers, TabPFN was pre-trained on a specific distribution of small synthetic tabular data. As a result, TabPFN only excels in handling data with up to 10,000 samples, 500 features, and 10 classes. Furthermore, TabPFN requires a GPU for inferences. For datasets with more samples or features, TabPFN requires more VRAM and, thus, a more expensive GPU. At the same time, even with a GPU, the inference speed may be slower than for traditional forensic genetics models. Lastly, TabPFN, even as a foundation model, might still benefit from integrating more domain expertise as well as advanced preprocessing, data cleaning, or feature engineering. TabPFN shall empower, not replace, data scientists in forensic genetics.

Our results emphasize limitations for classification: First, a small number of training samples as e.g. in Central Asia leads to poorer results. Especially, for the intra-European classification, this is a problem and leads even for TabPFN to a low accuracy. Hence, before using TabPFN to classify individuals into populations, researchers should apply cross validation to ensure that the performance of the classifier is good enough for their research question. Ideally, if there is uncertainty in the choice of populations, one should consider either combining or separating them. Second, we have only studied classification into existing populations, which ignores the possibility of admixed individuals. The latter require estimation of admixture proportions, a task which can only be handled by the Admixture Model out of the box.

However, our results show that TabPFN outperforms all other, commonly used classifiers in the forensic genetics. Hence, we advocate the use of TabPFN to classify individuals into populations. Therefore, we provide an offline interface with some example data. However, the user is free to use the method on any other data.

Improving the performance of classification in BGA suggests that similar tasks can be improved as well. In particular, researchers can apply TabPFN to external visible characteristics, where eye color (Liu et al., 2009; Walsh et al., 2011) or the hair color prediction (Walsh et al., 2013) are considered. Moreover, we used TabPFN without optimizing (i) the input for the markers used or (ii) the weights of the neural network for the specific task. These two possibilities might even further increase the accuracy, although they come with a large computational cost. Since feature selection typically improves the performance of a classifier (Venkatesh and Anuradha, 2019), the accuracy of TabPFN might increase through choosing a marker set optimized for the considered populations, e.g., a marker set for intra-european classification.

## A Number of Individuals per Population

We present the number of individuals for every population, completing the information in Figure 1.

**Table S1.** Sample sizes on the continental level.

| Population | AFR | SAS | EUR | EAS | CAS | NAF | OCE | AMR | MEA |
| --- | --- | --- | --- | --- | --- | --- | --- | --- | --- |
| Number of Individuals | 827 | 729 | 830 | 935 | 64 | 29 | 77 | 473 | 377 |

**Table S2.** Sample sizes on the intra-continental level.

| Population | TSI | BAS | SAR | IBS | CEU | FIN | GBR | FRA | TUR | RUS |
| --- | --- | --- | --- | --- | --- | --- | --- | --- | --- | --- |
| Number of Individuals | 107 | 23 | 28 | 107 | 99 | 99 | 91 | 28 | 28 | 25 |

## B Exact Values for the ROC AUC, Accuracy and LogLoss

We present the rounded values for the mean of the three considered evaluation metrics; compare with Figure 2. Recall that classification problem (1) is continental with 8 populations, while problem (2) is inter-European with 10 populations.

**Table S3.** ROC AUC for the different classification problems (1) and (2).

| Classifier | TabPFN | PLS-DA | SNIPPER | AM | XGBoost | RF |
| --- | --- | --- | --- | --- | --- | --- |
| ROC AUC Problem (1) | 1 | 0.96 | 0.86 | 0.96 | 0.94 | 0.99 |
| ROC AUC Problem (2) | 0.84 | 0.78 | 0.67 | 0.73 | 0.79 | 0.79 |

**Table S4.** Accuracy for the different classification problems (1) and (2).

| Classifier | TabPFN | PLS-DA | SNIPPER | AM | XGBoost | RF |
| --- | --- | --- | --- | --- | --- | --- |
| Accuracy Problem (1) | 0.93 | 0.86 | 0.84 | 0.82 | 0.91 | 0.90 |
| Accuracy Problem (2) | 0.48 | 0.47 | 0.43 | 0.40 | 0.42 | 0.45 |

**Table S5.** Log Loss for the different classification problems (1) and (2).

| Classifier | TabPFN | PLS-DA | SNIPPER | AM | XGBoost | RF |
| --- | --- | --- | --- | --- | --- | --- |
| Log Loss Problem (1) | 0.20 | 0.71 | 2.96 | 2.27 | 0.30 | 0.45 |
| Log Loss Problem (2) | 1.37 | 3.08 | 3.01 | 13.17 | 1.84 | 1.74 |

## C Results for XGBoost and Random Forest

We present the confusion matrices for XGBoost and Random Forest on both, the continental as well as the intra-European level; see Figures 3 and 4 for the results using TabPFN, PLS-DA, SNIPPER and the Admixture Model.

**Figure S1.**
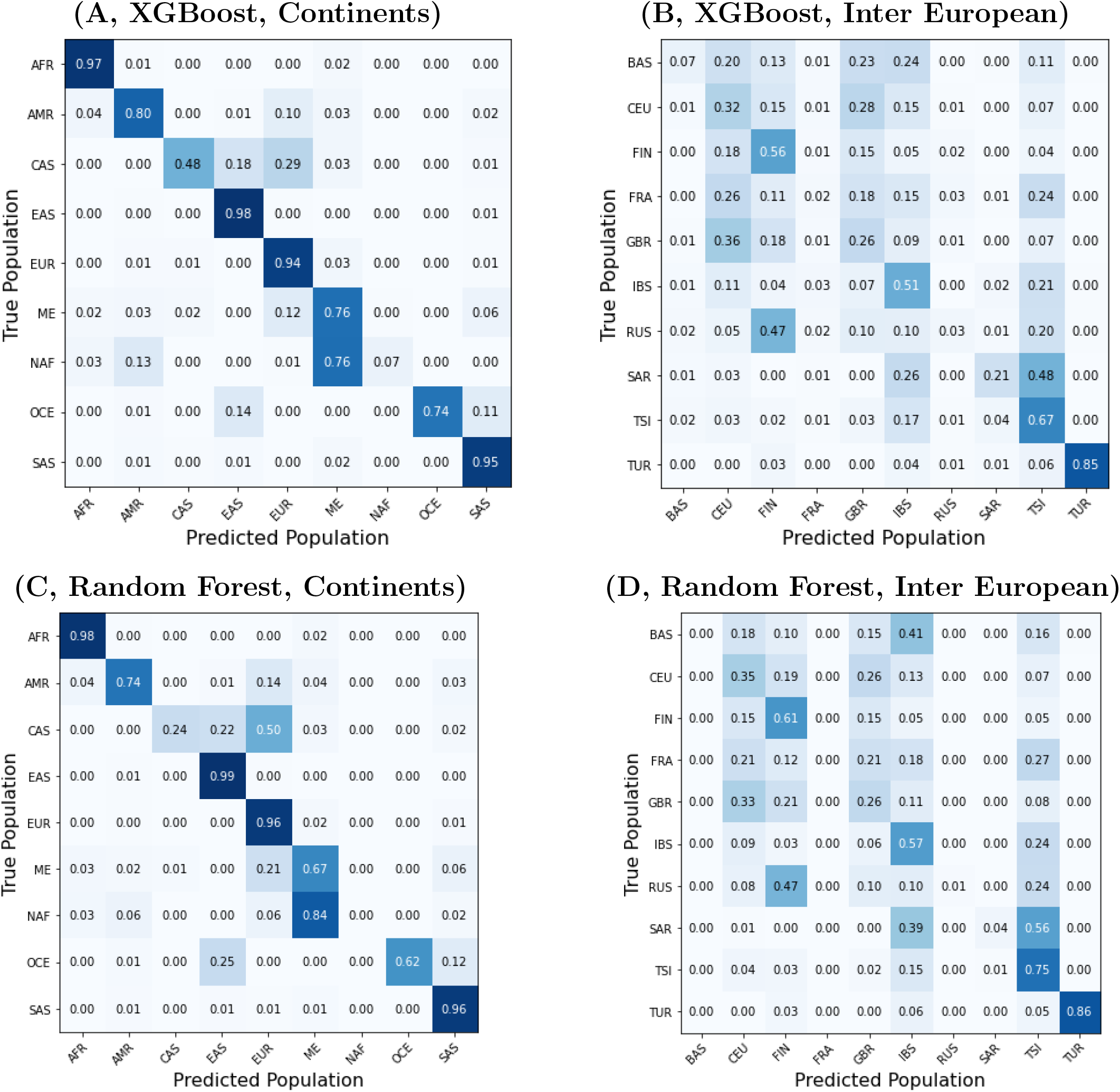
Confusion matrices for intra-European classification (case 2). All rows sum to 1 up to a rounding error. (A) XGBoost, Continents; (B) XGBoost, Inter European; (C) Random Forest, Continents; (D) Random Forest, Inter European

## Footnotes

1 https://github.com/PriorLabs/tabpfn

1 We employ scikit-learn’s one-vs-rest method to extend ROC AUC to multiclass.

